# *Phytophthora capsici* carries and differentially expresses genes that encode key enzymes for the synthesis, transport and processing of small RNAs

**DOI:** 10.1101/2025.09.21.677644

**Authors:** Jacobo Sevillano-Serrano, Fernando Uriel Rojas-Rojas, Alfonso Méndez-Bravo, Nancy Calderón-Cortés, Harumi Shimada-Beltrán, Julio C. Vega-Arreguín

## Abstract

The small RNA (sRNA) pathway is an epigenetic mechanism that has recently gained attention due to its suggested role in regulating virulence of plant pathogens. This gene silencing process has been observed in certain species of the oomycete genus *Phytophthora*. However, little is known about this pathway in *Phytophthora capsici*, a pathogen with a broad host range that affects many important food crops. In the present study, using bioinformatics approaches on the reference genome, transcriptome, and proteome of *P. capsici*, we identified and analyzed key genes and proteins involved in the synthesis, transport, and processing of sRNAs.

Our results showed that the *P. capsici* genome encodes a DCLα, DCLβ, exportin-5A, RDR, and six AGO proteins, suggesting the presence of a complete sRNA pathway in this pathogen. These genes were syntenic, structurally similar, and phylogenetically related to other oomycetes of the genus *Phytophthora*. We also analyzed their expression levels after infecting chili pepper and broccoli across two generations, revealing different expression patterns depending on the pathogen infection history.

To our knowledge, this is the first report showing the presence of the exportin-5A gene in *P. capsici* and other oomycetes. Additionally, the expression of all these sRNA-related genes in the pathogen isolated from different hosts suggests that the host may influence the epigenetic regulation of *P. capsici* via the sRNA pathway. This study paves the way for functional studies to confirm the role of the sRNA pathway in regulating virulence in *P. capsici*.

## Introduction

*Phytophthora capsici* is a plant pathogenic oomycete [1–4] that belongs to the Chromista kingdom and that threatens agricultural and natural ecosystems [5–7]. Although oomycetes resemble filamentous fungi, they are phylogenetically related to aquatic organisms such as diatoms and algae [8–11]. *P. capsici* can infect a wide range of hosts in laboratory, greenhouse, and field conditions, affecting at least 26 plant families, including both ornamental and native plants [2]. This pathogen poses a significant threat to food security [4, 10, 12] because it causes diseases in various crops worldwide, such as chili peppers, cucumbers, watermelons, and squash, among many others [13–18]. *P. capsici* is a hemibiotrophic, soil-borne pathogen that exhibits mixed reproduction [19, 20], capable of infecting host plants at any growth stage, leading to seedling death, crown rot, leaf blight, and fruit rot [13, 21, 22]. This makes it an excellent model organism for studying the adaptation of plant pathogens to diverse hosts, including epigenetic mechanisms of gene silencing that could play a role in regulating this process [20].

Gene silencing mediated by small RNAs (sRNAs) is known to occur in oomycetes, including *P. infestans*, *P. sojae,* and *P. parasitica* [23–26], but the molecular mechanisms underlying this process are not well characterized in these organisms [27]. sRNAs are short RNA sequences, 20 to 40 nucleotides (nt) long, that do not encode proteins but are known to play key roles in various eukaryotic pathogens, including the regulation of endogenous biological processes and interactions with hosts [28–33]. In general, sRNAs are classified as microRNAs (miRNAs), small interfering RNAs (siRNAs), and small piwi-associated RNAs (piRNAs) [34]. In plants and oomycetes, sRNAs are synthesized and processed by the action of Dicer-like enzymes (DCL) [35], whereas in animals and mammals, this occurs via Drosha and Dicer enzymes [36,37]. sRNAs are transported from the nucleus to the cytoplasm via exportin-5/HASTY [38,39], where they bind to an enzyme of the Argonaute (AGO) family to form the RNA-induced silencing complex (RISC). This complex binds to a target messenger RNA (mRNA) through base complementarity between the sRNA and mRNA [46]. The AGO enzyme then performs its ribonuclease activity and cleaves the target mRNA. This results in the regulation of the target mRNA at the post-transcriptional level, a process known as post-transcriptional gene silencing (PTGS). However, miRNA regulation can occur before transcription of the target mRNA, resulting in transcriptional gene silencing (TGS) [47–49]. Subsequently, the RNA-dependent RNA polymerase (RDR) may recognize the cleaved mRNA fragments and synthesize double-stranded RNAs (dsRNAs). These can be incorporated into the DCL enzyme, generating small secondary RNAs that amplify the silencing signal for a particular gene by binding to AGO enzymes and recognizing complementary sequences [43,44]. The understanding of how this gene silencing process by sRNAs, along with the enzymes involved, regulates pathogenicity and development in *Phytophthora* species is limited [50]. Indeed, the few reports on epigenetic regulation in pathogenic oomycetes have primarily focused on histone methylation, acetylation, and deacetylation, as well as small RNAs related to effector genes [51–54].

In this study, we aimed to identify, characterize *in silico*, and evaluate the expression of key genes encoding enzymes associated with the sRNA pathway in *P. capsici,* and to determine their expression patterns in the pathogen after infection of different host plants. This analysis could support future functional studies to describe the specific roles of each enzyme in the sRNA pathway in *P. capsici* and their influence on regulating pathogenicity and growth.

## Materials and methods

### Identification and phylogenetic analysis of sRNA-related enzymes in *P. capsici*

DCLs, exportin-5 (Exp5A), AGO, and RDR protein sequences were searched in the *P. capsici* LT1534 proteome (GCA_000325885.1) from the PhycoCosm portal of the Joint Genome Institute (JGI) [20] using Hidden Markov Models (HMM) based on the seed alignments of functional domains of each enzyme family obtained from the Pfam v35.0 database [55].

The HMM probability distribution of the characteristic domains for each enzyme was built using the HMMER v3.3.2 program [56], and proteins containing the specified number and position of characteristic domains within each protein family were identified. Proteins with two RNAase III domains at the C-terminus were classified as DCL enzymes. Sequences containing PAZ, MID, and PIWI domains were classified as AGO proteins. To be considered RDRs, sequences must contain an RdRP domain. Sequences lacking a domain were considered pseudogenes/proteins.

The conserved domains Xpo1 and exportin-5 from the *Xenopus tropicalis* [57] exportin-5 sequence were used to construct an HMM with HMMER c3.3.2 [55,56] to identify the Exp5A protein in *P. capsici*.

For phylogenetic analysis, homologous protein sequences reported in other oomycetes were retrieved from the NCBI portal. Homologous protein sequences from *Arabidopsis thaliana*, *H. sapiens*, and *Mus musculus* were used as outgroups. Subsequently, multiple sequence alignments for each protein family were performed using MUSCLE in MEGA11 [58], followed by sequence curation with GBlocks v0.91b [59]. Phylogenetic reconstruction was then performed using maximum likelihood (ML) with the Akaike Information Criterion (AIC) in PhyML [60], selecting the best-fitting substitution model, and supporting branches with 100 bootstrap iterations. Tree topology visualization was performed using TreeDyn v1.98.3 and Evolview [61,62].

Characteristic domains were constructed using the NCBI Conserved Domain Search tool [63] and visualized with TBtools-II v1.120 [64]. The AGO nomenclature reported for other oomycetes has been retained. However, letters of the alphabet were assigned to distinguish proteins that were not reported in other pathogens. Finally, the subcellular localization of the genes was predicted using Predict Protein LocTree3v4.0 [65].

### Characteristics of exportin-5A of *P. capsici*

Since no genes or proteins involved in sRNA export have been identified in *Phytophthora* species, we analyzed the genes and proteins found in *P. capsici* that have sRNA export features similar to those of exportin-5 genes reported in other organisms. The genetic neighborhood of the *P. capsici* Exp5A gene was examined using the genome browser tool at the Joint Genome Institute (JGI) [66] and was compared with the genomes of *P. infestans*, *P. sojae*, and *P. ramorum*. The features of *P. capsici* scaffolds were visualized using the synteny tool on the JGI website [66]. Multiple sequence alignment of Exp5A genes from *Phytophthora*, *H. sapiens*, and *M. musculus* was performed using EMBOSS Needle [67] and MUSCLE [58] to identify the conserved domains IBN_N, Xpo1, and exportin-5. The abundance of amino acids at each position in the primary sequence of these proteins was analyzed using the Weblogo program [68].

### Synteny analysis and gene interaction networks

To understand the conservation of genes encoding key enzymes for the biogenesis and processing of sRNAs in *P. capsici*, we performed a synteny and collinearity analysis using the Synteny OneStep MCScanX tool [73] in TBtools-II v1.120 [64], across the genomes of *P. capsici* [69], *P. infestans* [70], *P. sojae* [71], and *P. ramorum* [72], obtained from the NCBI and JGI databases.

To predict whether the identified sRNA pathway proteins of *P. capsici* could interact with each other or with other related proteins, we performed *in silico* protein-protein interaction analysis. Protein-protein interaction networks of the DCL, Exp5A, AGO, and RDR genes were built in the STRING v11.5 database [74], with a medium confidence interaction score (0.40) and 1st and 2nd shell no more than five interactions. The confidence score selected indicates a medium level of confidence in the predicted protein-protein interaction or biological significance. The 1st and 2nd shell parameter “no more than five interactions”, specify that only the top five most likely interactions between the proteins being analyzed and another set of the top five related proteins are displayed.

### Influence of hosts on the expression of *P. capsici* sRNA pathway-related genes

#### Infection and isolation of *P. capsici* from plant tissue

To determine the expression pattern of sRNA pathway-related genes in *P. capsici* after infecting different hosts, leaves from chili and broccoli were collected from plants grown in greenhouse conditions for 4-6 weeks and infected with agar plugs containing fresh mycelium of *P. capsici* D3 as described previously [75]. Infection assays were performed as follows: the detached leaves were washed with 70% ethanol, rinsed three times with sterile water, and then placed in a sterile humidity chamber with the underside facing up. Next, a ∼2 mm wound was made in the center of each leaf and inoculated with a 5 mm diameter plug of 8-day-old mycelium of *P. capsici* D3. The leaves were incubated at 28°C in the dark, and infection was monitored at 24, 48, and 72 hours post-inoculation (hpi). Each experiment was performed six times using four leaves per plant species. The infected area on each leaf was measured using ImageJ software [76], and statistical analysis, including normality tests and ANOVA (p<0.05), were performed.

*P. capsici* D3 was isolated from infected tissue of chili and broccoli (primary infections, I-chili and I-broccoli) at 72 hpi and grown in the dark at 28°C in Petri dishes with V8 medium [77] supplemented with the antimicrobials benomyl (100 mg/L), rifampicin (80 mg/L), ampicillin (100 mg/L), streptomycin (50 mg/L), and kanamycin (50 mg/L). These first isolates of *P. capsici* D3 (Pc-chili and Pc-broccoli) were used to reinfect new chili plant tissue (secondary infections, I-ch-ch and I-br-ch). The isolates from these secondary infections (Pc-ch-ch and Pc-br-ch) were cultured in the same way as those from the primary infections. Both primary and secondary isolates of *P. capsici* D3, obtained after infecting the two hosts, were grown for 8 days, and the mycelia were collected for total RNA extraction, which was then used in gene expression assays.

#### RT-qPCR assays

Relative gene expression in the mycelia of the parental *P. capsici* D3 without infection history (control), primary (Pc-chili, Pc-broccoli), and secondary isolates (Pc-ch-ch, Pc-br-ch), were quantified by RT-qPCR. Primers designed and used for each gene are listed in S3 Table.

To determine the expression of DCLs, Exp5A, AGO, and RDR genes of *P. capsici,* primers were designed using the NCBI Primer-BLAST program [78] and validated by end-point PCR with a temperature gradient, using *P. capsici* D3 genomic DNA obtained with the Plant/seed DNA Miniprep kit (ZYMO RESEARCH) following the manufacturer’s instructions.

Gene expression was analyzed by quantitative PCR (qPCR) using cDNA libraries prepared through reverse transcription (RT) from total RNA of five *P. capsici* mycelium samples (D3, Pc-chili, Pc-broccoli, Pc-ch-ch, and Pc-br-ch) obtained as described above. Total RNA was extracted using the Quick-RNA Plant Miniprep kit (Zymo Research) according to the manufacturer’s guidelines, and its integrity was assessed by agarose gel electrophoresis and NanoDrop. cDNA synthesis was performed using the Revert First Strand cDNA Synthesis kit (Thermo Scientific) by retro-transcription reactions of 20 µL containing 4 µL 5X Buffer, 2 µL dNTPs (10 mM), 0.5 µL RiboLock, 1 µL oligo(dT)_18_, 10 units of RevertAid Reverse Transcriptase, 1000 ng of total RNA, and nuclease-free water in a thermal cycler with the following program: 42 °C for 60 min, 70 °C for 5 min, and hold at 4 °C.

qPCR was performed in 10 µL reactions consisting of 5 µL SYBR Green Real-Time PCR Master Mix 2X (Applied Biosystems | Thermo Fisher Scientific), 1 µL Forward Primer (2 mM), 1 µL Reverse Primer (2 mM), 1 µL cDNA (50 ng/µL), and 2 µL nuclease-free water on the StepOne Real-Time PCR System (Applied Biosystems). The qPCR conditions included an initial denaturation at 95°C for 10 min, followed by 40 cycles of 95°C for 15 s, then 54-68 °C for 30 s, and 72°C for 30 s; a third step involved increasing the temperature from 65°C by 0.3 °C every 0.1 min up to 95°C, and a final step at 95°C for 15 s. The 2ΔΔCt method [79] was used to determine the relative expression levels of each gene, with elongation factor 1α as the endogenous control gene. The assay was conducted in triplicate with two biological replicates. Graphs displaying gene expression levels, validated by ANOVA statistical analysis (p<0.05), were generated using GraphPad Prism7 [80].

## Results

### *P. capsici* encodes DCLs, Exp5A, AGOs, and RDR genes that are phylogenetically related to those in other pathogenic oomycetes

After the construction of HMMs and analysis of the *P. capsici* proteome, we identified key proteins related to sRNA biogenesis and processing. The characteristics of the genes encoding two DCLs (DCLα and DCLβ), one exportin 5 (Exp5A), six argonauts (A, B, C, D, E, and F), and one RDR are summarized in Table 1.

**Table 1.**
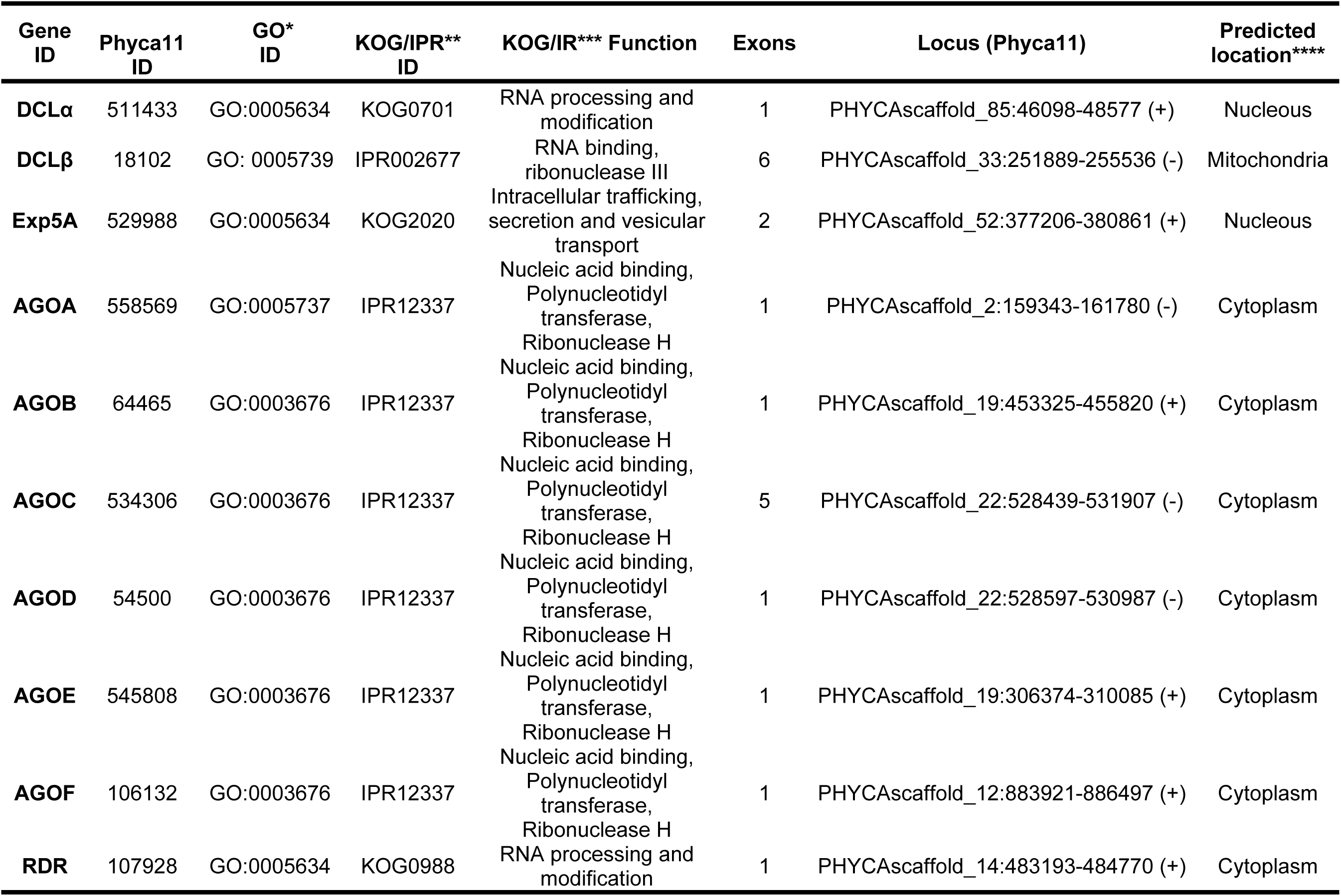
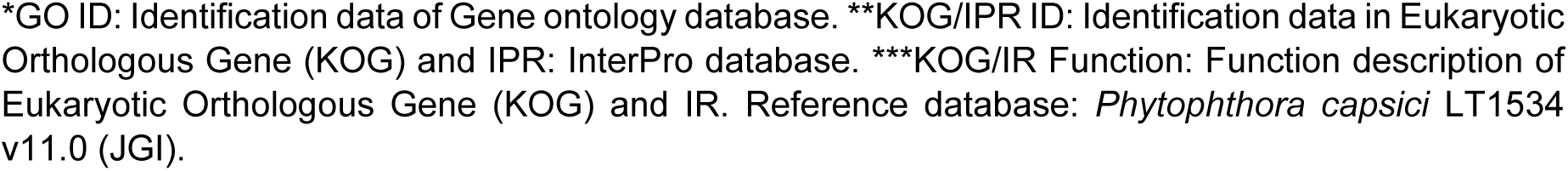
Characteristics of genes encoding key enzymes in the *P. capsici* sRNAs pathway.

| Gene ID | Phyca11 ID | GO* ID | KOG/IPR** ID | KOG/IR*** Function | Exons | Locus (Phyca11) | Predicted location**** |
| --- | --- | --- | --- | --- | --- | --- | --- |
| DCLα | 511433 | GO:0005634 | KOG0701 | RNA processing and modification | 1 | PHYCAscaffold_85:46098-48577 (+) | Nucleous |
| DCLβ | 18102 | GO: 0005739 | IPR002677 | RNA binding, ribonuclease III | 6 | PHYCAscaffold_33:251889-255536 (-) | Mitochondria |
| Exp5A | 529988 | GO:0005634 | KOG2020 | Intracellular trafficking, secretion and vesicular transport | 2 | PHYCAscaffold_52:377206-380861 (+) | Nucleous |
| AGOA | 558569 | GO:0005737 | IPR12337 | Nucleic acid binding, Polynucleotidyl transferase, Ribonuclease H | 1 | PHYCAscaffold_2:159343-161780 (-) | Cytoplasm |
| AGOB | 64465 | GO:0003676 | IPR12337 | Nucleic acid binding, Polynucleotidyl transferase, Ribonuclease H | 1 | PHYCAscaffold_19:453325-455820 (+) | Cytoplasm |
| AGOC | 534306 | GO:0003676 | IPR12337 | Nucleic acid binding, Polynucleotidyl transferase, Ribonuclease H | 5 | PHYCAscaffold_22:528439-531907 (-) | Cytoplasm |
| AGOD | 54500 | GO:0003676 | IPR12337 | Nucleic acid binding, Polynucleotidyl transferase, Ribonuclease H | 1 | PHYCAscaffold_22:528597-530987 (-) | Cytoplasm |
| AGOE | 545808 | GO:0003676 | IPR12337 | Nucleic acid binding, Polynucleotidyl transferase, Ribonuclease H | 1 | PHYCAscaffold_19:306374-310085 (+) | Cytoplasm |
| AGOF | 106132 | GO:0003676 | IPR12337 | Nucleic acid binding, Polynucleotidyl transferase, Ribonuclease H | 1 | PHYCAscaffold_12:883921-886497 (+) | Cytoplasm |
| RDR | 107928 | GO:0005634 | KOG0988 | RNA processing and modification | 1 | PHYCAscaffold_14:483193-484770 (+) | Cytoplasm |

\*GO ID: Identification data of Gene ontology database. \*\*KOG/IPR ID: Identification data in Eukaryotic Orthologous Gene (KOG) and IPR: InterPro database. \*\*\*KOG/IR Function: Function description of Eukaryotic Orthologous Gene (KOG) and IR. Reference database: *Phytophthora capsici* LT1534 v11.0 (JGI).

Phylogenetic analysis of the amino acid sequences of DCLα and DCLβ showed that both *P. capsici* proteins are closely related to the enzymes from *P. ramorum and P. sojae* (Fig 1A). DCLα was grouped into a clade with DCL1 proteins from *P. infestans*, *P. sojae*, *P. nicotianae,* and *P. ramorum*. Meanwhile, DCLβ was grouped with DCL2 from *P. infestans*, *P. sojae,* and *P. ramorum,* as well as the Drosha proteins from *D. melanogaster* and *H. sapiens*. Both DCLα and DCLβ possess characteristic domains of the Dicer-like family (Fig 1A).

**Fig 1.**
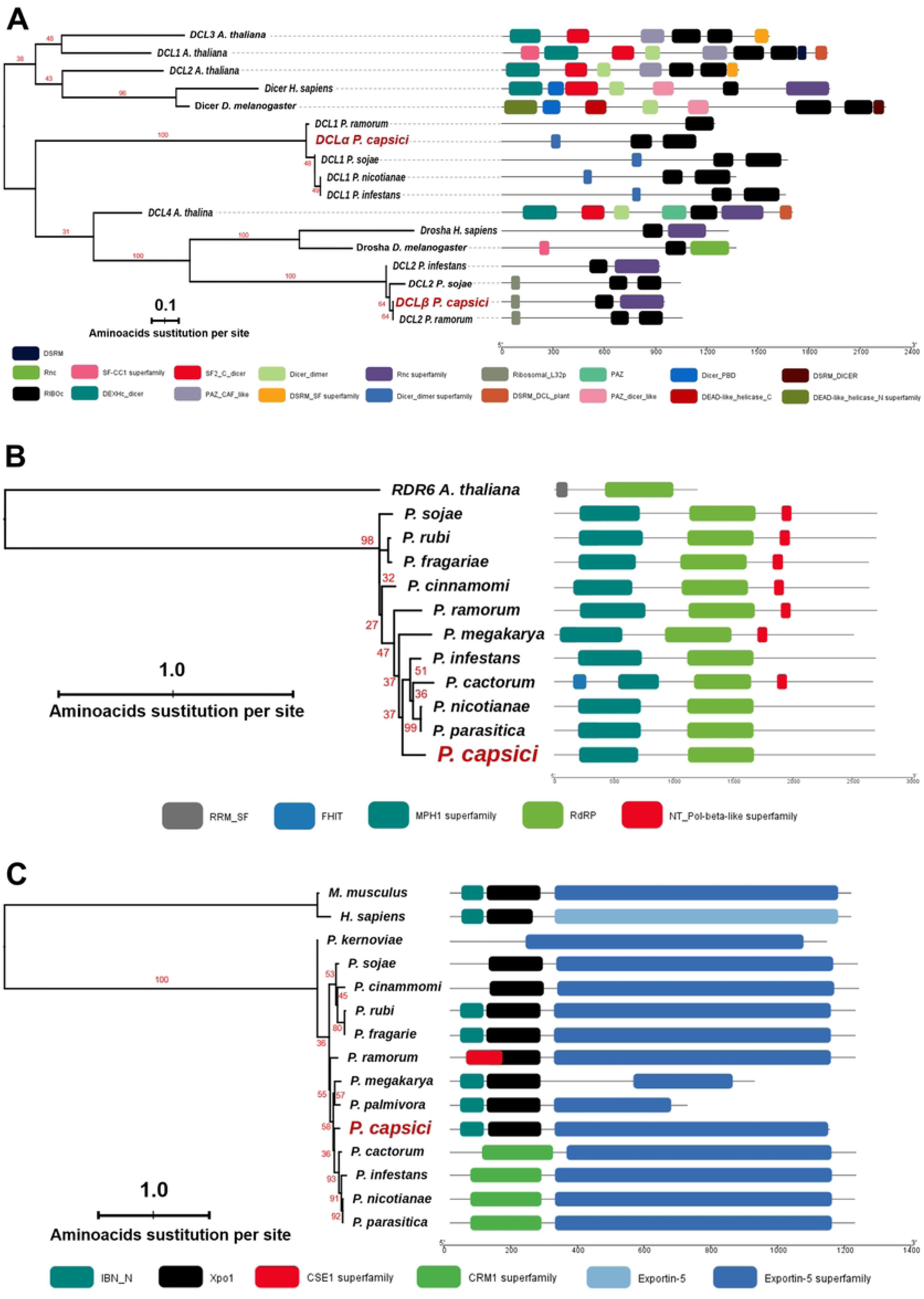
Phylogenetic reconstruction of DCL, RDR, and exportin-5 in *Phytophthora* species. (A) Phylogenetic tree of Dicer-like proteins from *P. capsici* and other species. *A. thaliana* DCL proteins 1-4, and Dicer and Drosha from *H. sapiens* and *M. musculus* were used as outgroups. (B) Phylogenetic tree of RNA-dependent RNA polymerase of *P. capsici* with RDR from other oomycetes, using *A. thaliana* RDR6 protein as an outgroup. (C) Phylogenetic tree of exportin 5 proteins in *Phytophthora* species, with exportin-5 from *M. musculus* and *H. sapiens* used as outgroups. The conserved domains of the proteins are highlighted in color on the right side of each panel. The scale bar indicates the size of the protein in amino acids. Colored rectangles with rounded corners indicate the protein domains.

The RDR identified in *P. capsici* contains the key domains known for this protein (Fig. 1B) and is mainly related to those in *P. infestans*, *P. cactorum*, *P. nicotianae*, and *P. parasitica*. The topology of the RDR phylogeny shows that these protein sequences are more closely related among *Phytophthora* species than to RDR6 of *A. thaliana*. Additionally, no potential RDR-related genes were found in species such as *P. kernoviae* or *P. palmivora*.

The phylogenetic reconstruction of Exp5A with homologues from other *Phytophthora* species indicated that Exp5A of *P. capsici* presented a closer phylogenetic relationship with the homologue of *P. cactorum*, *P. infestans*, *P. nicotianae* and *P. parasitica* (Fig 1C). In general, the oomycete exportin-5 clustered distantly from *M. musculus* and *H. sapiens* homologues. All proteins of the *Phytophthora* species contained the IBN_N, Xpo1, and exportin-5 domains, which are crucial for the functioning of this family of enzymes.

The phylogenetic reconstruction of the six argonautes of *P. capsici* (A, B, C, D, E, and F) shows that these enzymes grouped into two main clades of homologous AGOs from oomycetes such as *P. infestans*, *P. sojae*, *P. cactorum*, *P. cinammomi*, *P. fragarie*, *P. kernoviae*, *P. megakarya*, *P. nicotianae*, *P. palmivora*, *P. parasitica*, *P. rubi* and *P. ramorum* (Fig 2).

**Fig 2.**
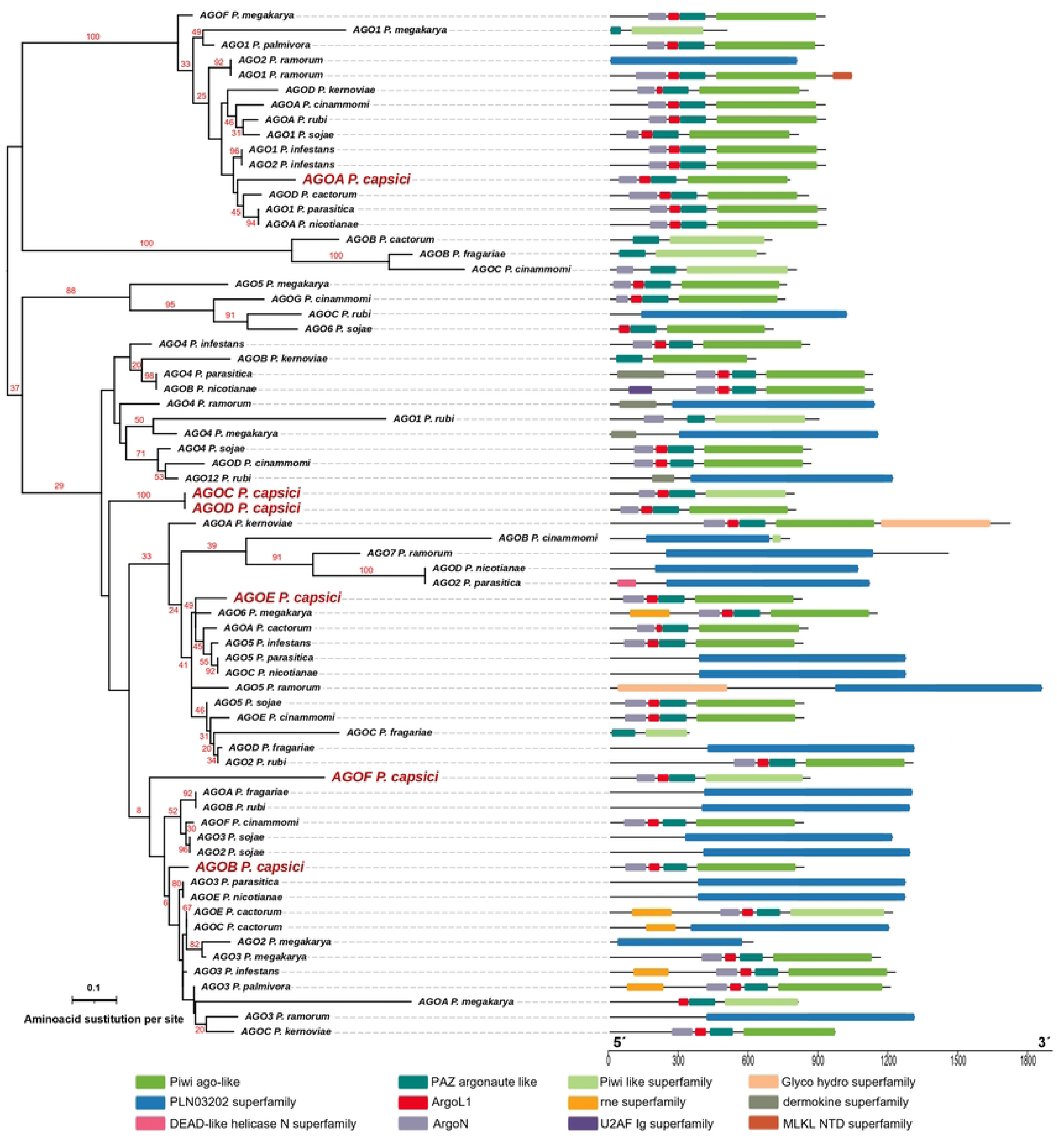
Phylogenetic reconstruction of AGO in *Phytophthora* species. Phylogenetic reconstruction of Argonaute proteins from *Phytophthora capsici* and other oomycetes. The scale bar indicates the size of the protein in amino acids. The conserved domains of the proteins are shown in color on the right side of the phylogeny.

AGOA is grouped into clade I with AGO1 from *P. parasitica*, *P. megakarya*, *P. palmivora*, *P. ramorum*, *P. sojae*, and *P. infestans*. In contrast, the other AGOs of *P. capsic*i fell into clade II. AGOB is mainly related to AGO3 from *P. parasitica* and AGOE from *P. nicotianae*. However, AGOC and D and F clustered in a separate clade, while AGOE shows a closer phylogenetic relationship to AGO6 from *P. megakarya*, AGOA from *P. cactorum*, AGO5 from *P. infestans*, AGO5 from *P. parasitica*, and AGOC from *P. nicotianae*. The primary structure of these proteins indicates that the *P. capsici* argonautes and the other AGO proteins analyzed vary in size and differ in the presence of characteristic domains of the argonaute proteins (ArgoN, AgoL, PAZ, and PIWI). Supporting Information 1 (S1 Table) lists the IDs and protein sequences used in the phylogenetic analyses.

The genomic neighborhood of the *P. capsici* DCLs, AGOs, and RDR genes showed that they were located on different scaffolds within gene-rich regions and some transposable element regions. However, some genes are situated relatively close to these sites (S1 Fig.). Additionally, the LocTree3v4.0 software indicated that DCLα and DCLβ proteins might be located in the nucleus and mitochondria, respectively, while AGOs and RDRs are likely located in the cytoplasm of cells (Table 1).

### *P. capsici* exportin-5A is located in gene-rich and conserved genomic regions

To determine the genomic neighborhood of the *P. capsici* Exp5A gene, we used the browser tool of the JGI (Fig 3A). We found that the gene was located on scaffold 52:377206-380861 (+) in *P. capsici* LT1534. Exp5A gene appears to be in a gene-rich region, with neighboring genes on the left and right corresponding to a drug/metabolite transporter (DMT) and a galactosyltransferase, respectively. Exp5A contains two exons and is highly similar to other homologous genes in the genomes of *P. sojae* (ID 465489, Physo3) and *P. ramorum* (ID 81240, *P. ramorum* v1.1).

**Fig 3.**
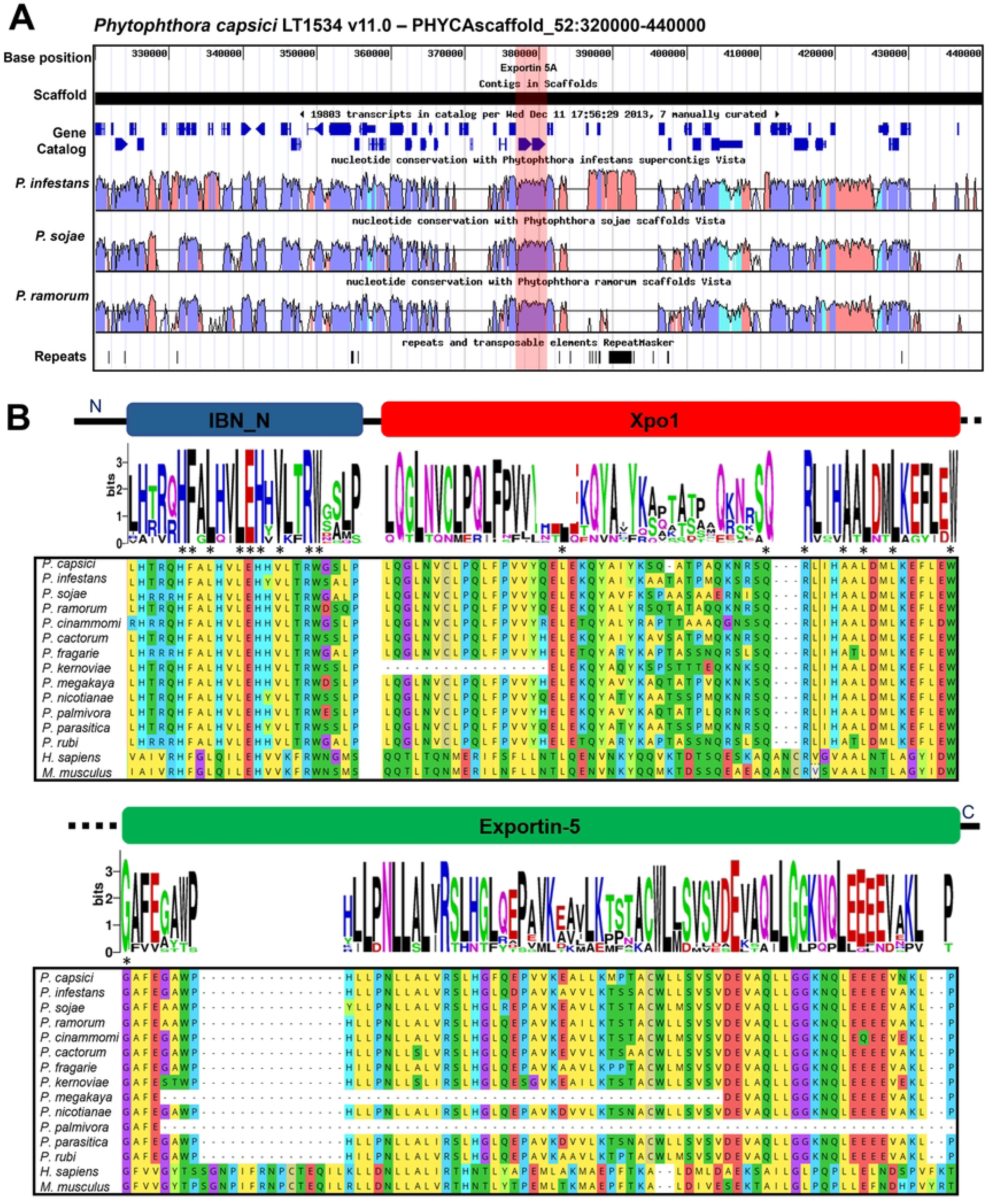
Genomic neighborhood and conserved domains of exportin-5. (A) Genomic neighborhood of *P. capsici* exportin-5A gene shown in red stripe, located in gene-dense region and highly conserved regions between *P. infestans*, *P. sojae* and *P. ramorum* genomes, conserved region cyan: UTR, blue: exons, red: introns. (B) Sequence alignment of exportin-5 proteins divided by the conserved domains of IBN_N, Xpo1 and Exportin-5 in various species of *Phytophthora*, *H. sapiens* and *M. musculus* linked to a weblogo plot of abundance of amino acids at a determined position. * Amino acid conservation among all ranked organisms. - Amino acid misalignment.

Global alignment of the *P. capsici* Exp5A protein with homologs from *M. musculus* and *Homo sapiens* showed 24.6% and 24.2% identity, and 41% and 41.5% similarity, respectively. However, among oomycetes such as *P. infestans*, *P. sojae*, and *P. ramorum*, the identity was 86.2%, 86.8%, and 87.2%, with 92.9%, 93%, and 92.8% similarity, respectively. Domain analysis of *P. capsici* Exp5A indicated that it contains the three characteristic domains of exportin-5: IBN_N, CRM1, Xpo1, and exportin-5.

Multiple alignment of oomycete exportin-5 proteins and their *H. sapiens* and *M. musculus* homologs showed decreased similarity and differences in the amino acid residue composition of the Xpo1 and exportin-5 domains (Fig. 3B). Likewise, *P. kernoviae* showed the loss of a large region of this domain. The exportin-5 domain was also found to be the most variable between oomycetes and mammals, and *P. megakarya* and *P. palmivora* presented a smaller region of the domain compared to the other organisms (Fig. 3B).

### Genes of the sRNA pathway of *P. capsici* revealed synteny with four oomycete genomes and the formation of protein interaction networks

The synteny analysis of the genomes of *P. capsici*, *P. infestans*, *P. sojae*, and *P. ramorum* indicated that the key genes in the sRNA pathway DCLα, DCLβ, Exp5A, AGOA, AGOB, and RDR are syntenic among the four genomes (Fig 4A and S2 Table). In contrast, the AGOC, AGOD, AGOE, and AGOF genes of *P. capsici* are not syntenic with those in the other oomycetes (*P. infestans*, *P. sojae*, and *P. ramorum*) (Fig 4A and S2 Table). Overall, it appears that the DCLs, Exp5A, AGOs, and RDR genes are dispersed throughout the *P. capsici* genome, similar to those in *P. infestans* and *P. sojae*; in contrast, in *P. ramorum*, these genes are located in close proximity to each other.

**Fig 4.**
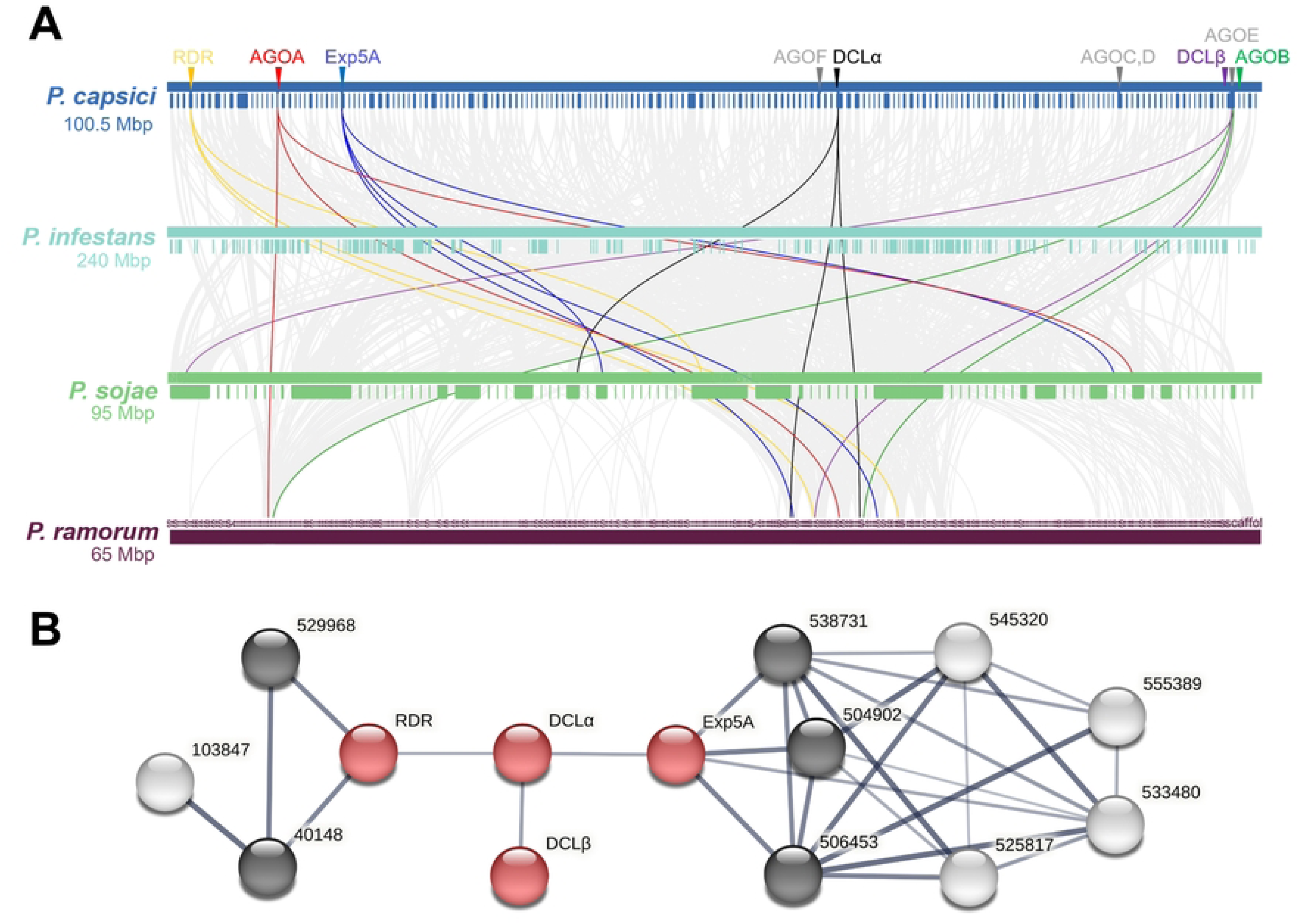
Synteny and protein-protein interaction networks of key genes in the small RNA pathway. (A) Synteny of key genes in the small RNA pathway in four oomycete species. Gray lines in the background indicate the collinear blocks within the *P. capsici* and other oomycete genomes, whereas the colored lines highlight the syntenic biosynthesis, transport and processing genes. (B) Protein-protein interaction networks of DCLα, DCLβ, Exp5A and RDR from *P. capsici*. Red nodes: interaction network of proteins related to sRNAs in *P. capsici*; black nodes: first shell of interactions; white nodes: second shell of interactions; shell number: proteome ID from the *P. capsici* v11 JGI database. Thicker dark line indicates higher score in protein-protein interaction. IDs: 504092 (GTPase Ran/TC4/GSP1 nuclear protein transport), 506453 (trafic intracellular, nuclear pore complex, Nup214/CAN component), 538731 (Ran GTPase-activating protein, RNA processing and modification), 525817 (ubiquitin-protein ligase), 545320 (karyopherin importin beta 1, intracellular trafficking, secretion and vesicular transport), 555389 (nuclear export signal-RNA export factor), 533480 (nuclear porin, structural constituent of nuclear pore), and 103847 (DNA excision repair protein XPA/XPAC/RAD14).

Predictive analysis of the protein-protein interaction networks of DCLα, DCLβ, Exp5A, and RDR (Fig 4B) suggests that they may interact within the same regulatory pathway with a mean confidence score of 0.4. Additionally, Exp5A potentially interacts with proteins 504092 (GTPase Ran/TC4/GSP1, nuclear protein transport), 506453 (traficc intracellular, nuclear pore complex, Nup214/CAN component), and 538731 (Ran GTPase-activating protein, RNA processing and modification) in 1st shell. There is also predictive evidence that it interacts with 525817 (ubiquitin-protein ligase), 545320 (karyopherin importin beta 1, intracellular trafficking, secretion, and vesicular transport), 555389 (nuclear export signal-RNA export factor), and 533480 (nuclear porin, structural constituent of nuclear pore) in 2nd shell. RDR potentially interacts with proteins 529968 (E3 ubiquitin ligase) and 40148 (structure-specific endonuclease ERCC1-XPF, ERCC1 complex) in the 1st shell, and with 103847 (DNA excision repair protein XPA/XPAC/RAD14) in the 2nd shell.

### *P. capsici* showed changes in virulence and different expression patterns of sRNA pathway genes after infecting chilli pepper and broccoli for two generations

To determine whether sRNA pathway genes are expressed in *P. capsici* after infecting different plant species with different infection histories over two generations, we analyzed their expression levels using RT-qPCR. First, we evaluated the development of *P. capsici* infections in chili pepper and broccoli leaves. The pathogen showed a specific infective ability on chili pepper and broccoli leaves with a progressive increase in the infected area at 24, 48, and 72 hpi (Fig 5A). Chili pepper leaves were more susceptible to infection by the pathogen at 72 hpi. In contrast, the pathogen exhibited limited infective capacity in broccoli leaves, indicating that this plant species was the most resistant to the pathogen. At 48 hpi, during secondary infections, *P. capsici* showed a statistically significant decrease in virulence (I-ch-ch and I-br-ch). However, at 72 hpi, no significant changes in virulence were observed under either condition.

**Fig 5.**
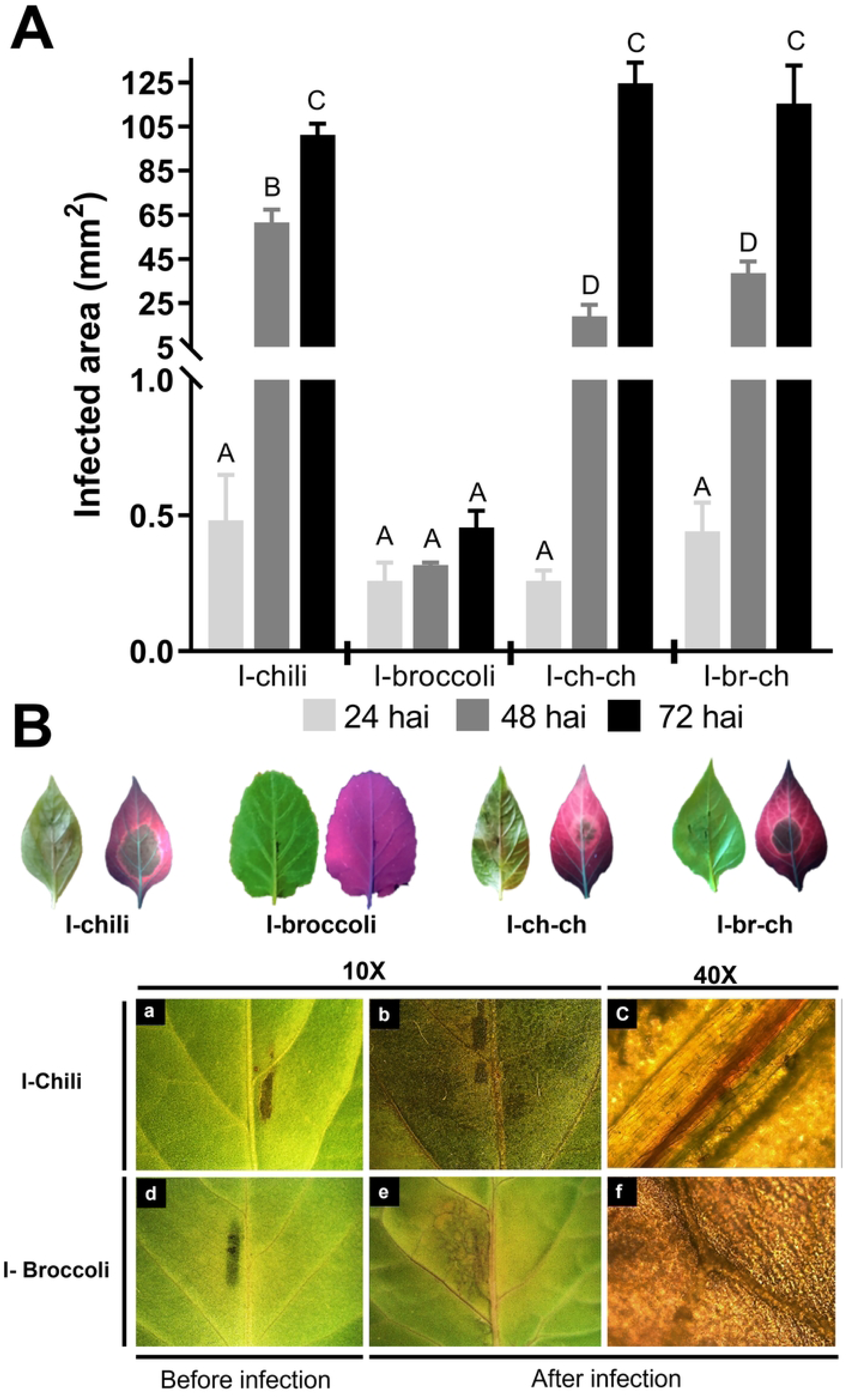
Infection caused by *P. capsici.* (A) Infection caused by *P. capsici* D3 in chili and broccoli leaves at 24, 48 and 72 hours post-inoculation (hpi). The bars indicate the standard error of the mean (SEM) of the infected area in mm^2^ from five independent experiments. Groups A, B, C, and D indicate ANOVA significant differences (p<0.05) in the development of infection within each plant species. (B) Phenotype of *P. capsici* infection in chili and broccoli. Left leaf of each host: visualization in natural light; the dark regions show the infected area. Right leaf: visualization with UV light (340 nm); the red regions indicate healthy tissue, the dark regions show the infected area in necrosis, and the orange regions show infection development in the biotrophic phase. Infection zoom: before and after infection (72 hpi) at 10X and 40X.

Infection in chili pepper leaves was characterized by necrosis in the center and along the veins as dark areas, probable biotrophy at the leaf edges and orange coloration observed under UV light. However, the leaves did not show chlorosis or wilting (Fig 5B). Meanwhile, infections in broccoli leaves showed necrosis around the inoculation site, followed by a chlorotic halo, probable biotrophy around the inoculation site and before the chlorotic area, and rippling at the leaf edges (Fig. 5B).

Subsequently, we identified a unique expression profile for each gene of *P. capsici* across the four experimental conditions (Pc-chili, Pc-broccoli, Pc-ch-ch, and Pc-br-ch) and the control D3 (S4 Table). The highest expression of DCLα was observed in the *P. capsici* D3 parental strain (control), which was significantly higher than that in the isolates from primary (Pc-chili, Pc-broccoli) and secondary (Pc-ch-ch, Pc-br-ch) infections (Fig 6A). However, Pc-ch-ch exhibited the lowest gene expression levels among the isolates. Additionally, the expression level of DCLβ indicated that in Pc-chili, Pc-broccoli (primary isolates), and Pc-ch-ch, Pc-br-ch (secondary isolates), gene expression was significantly reduced relative to D3 (Fig 6B). Nonetheless, Pc-broccoli and Pc-br-ch had significantly higher expression levels than Pc-chili and Pc-ch-ch, respectively.

**Fig 6.**
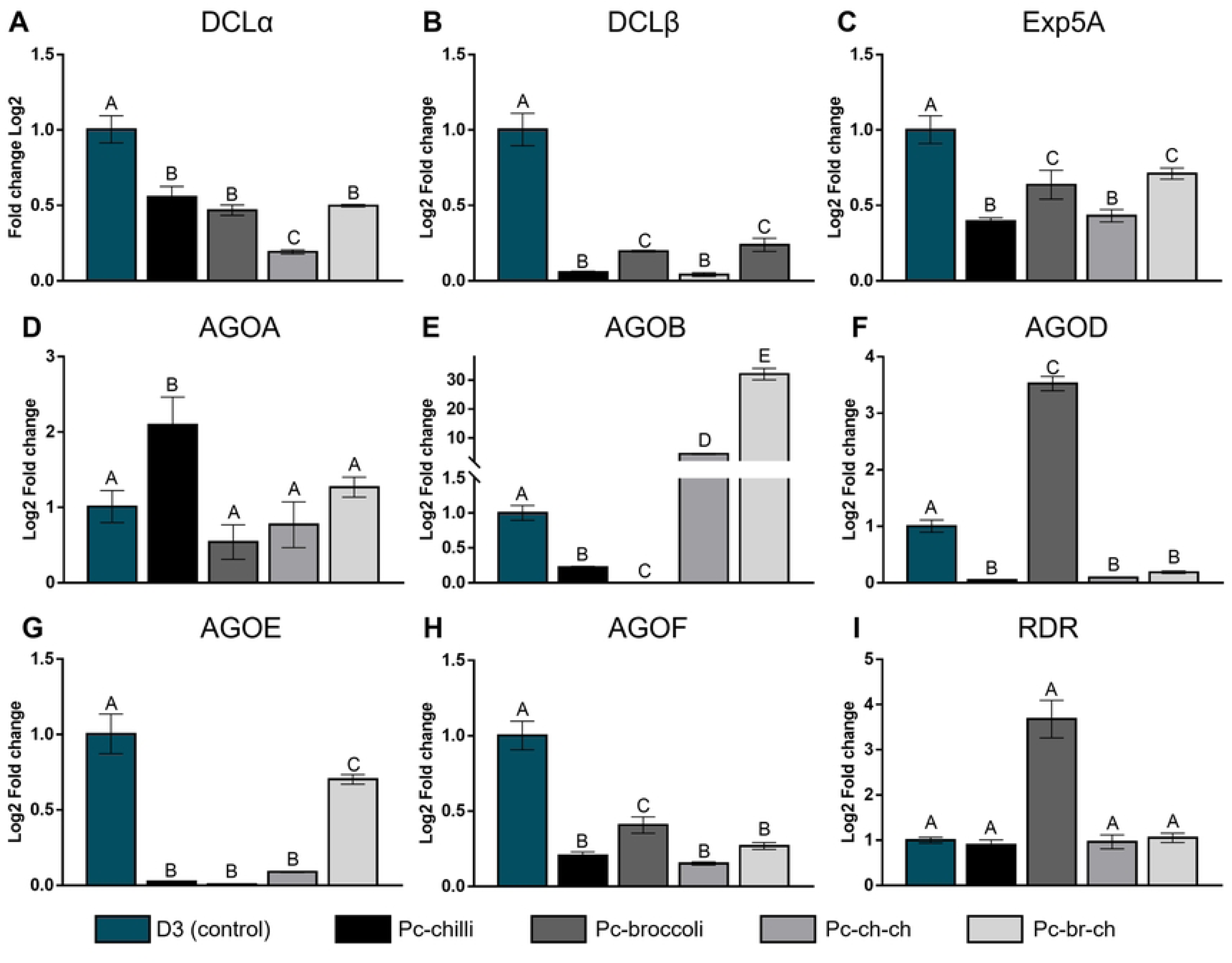
Relative expression of key genes in the small RNAs pathway in *P. capsici*. **(A)** DCLα. (B) DCLβ. (C) Exp5A. (D) AGOA. (E) AGOB. (F) AGOD. (G) AGOE. (H) AGOF and (I) RDR by 2ΔCt (Log2 fold change) from the mycelium of *P. capsici* D3 (control), Pc-chili and Pc-broccoli, Pc-ch-ch and Pc-br-ch, corresponding to two biological replicates in triplicate. The bars represent the standard error. Groups A, B, C, D, and E indicate significant ANOVA differences (p<0.05) between treatments for the same gene; EF-1α was used as an endogenous gene.

On the other hand, the relative expression level of Exp5A in *P. capsici* (Fig 6C) showed a pattern similar to that of DCLβ; that is, in D3, the expression of the gene was significantly higher than that in isolates from primary and secondary infections (Pc-chili, Pc-broccoli, Pc-ch-ch, and Pc-br-ch). Additionally, the expression level of Exp5A was significantly higher in Pc-broccoli and Pc-br-ch than in Pc-chili and Pc-ch-ch, with no significant differences between Pc-broccoli and Pc-br-ch, or between Pc-chili and Pc-ch-ch. When analyzing the relative expression profile of AGOA (Fig 6D), it was found that Pc-chili expressed the gene significantly higher than D3, Pc-broccoli, Pc-ch-ch, and Pc-br-ch, with no significant differences among the latter.

Meanwhile, the AGOB expression profile (Fig 6E) varied significantly across the treatments according to ANOVA analysis. Specifically, in Pc-chili and Pc-broccoli, AGOB expression was lower than that in the control, with no gene expression detected in Pc-broccoli. In contrast, AGOB was expressed at significantly higher levels in Pc-ch-ch and Pc-br-ch than in D3, showing a progressive increase in expression. The expression levels of AGOB were up to 3.5 times higher in Pc-ch-ch and 33 times higher in Pc-br-ch than in D3. No evidence of relative AGOC expression was observed. Conversely, the AGOD expression profile (Fig 6F) showed a significant increase in relative expression in Pc-broccoli compared to D3 (control). However, Pc-chili, Pc-ch-ch, and Pc-br-ch were expressed significantly less than the control, with no significant differences between them.

Our results also showed that the relative expression level of AGOE was higher in D3 than in Pc-chili, Pc-broccoli, Pc-ch-ch, and Pc-br-ch (Fig 6G). In contrast, the expression levels of the latter were higher than those of Pc-chili, Pc-broccoli, and Pc-ch-ch, with no significant differences among them, whereas the expression of AGOF decreased in the four evaluated treatments (Fig. 6H). However, Pc-broccoli expression was higher than that of Pc-chili, Pc-ch-ch, and Pc-br-ch, with no significant differences in AGOF expression in the latter. In contrast, the relative expression profile of the RDR gene (Fig 6I) showed similar levels of expression among the control D3, Pc-chili, Pc-ch-ch, and Pc-br-ch, but it was significantly higher in Pc-broccoli.

## Discussion

Epigenetic regulation mechanisms based on sRNAs allow for precise, dynamic, and reversible control of gene expression without changing the DNA sequence. They are essential for living [102] beings as they influence development [103], responses to biotic and abiotic stress [104], antiviral defense [105], silencing of transposable elements [106], and evolution [107].

Recently, the role of sRNAs in regulating oomycete infections has gained attention [26,28,53]. Therefore, understanding the molecular basis of sRNA synthesis, transport, and processing is essential for clarifying how virulence is regulated in plant pathogens, such as *P. capsici* [20,81]. In this study, we aimed to identify the genes and protein sequences of key enzymes involved in the sRNA pathway in *P. capsici* and analyze their relative gene expression after infection in different plant species.

Overall, our results revealed the presence of genes encoding key enzymes involved in the sRNA regulatory pathways in the *P. capsici* genome, including dicer-like (DCL), exportin (Exp), argonaute (AGO), and RNA-dependent RNA polymerase (RDR) genes. These sequences encode the characteristic functional domains associated with sRNA activity for each class, the presence of which in the *P. capsici* genome was mostly unknown. Specifically, we identified two dicer-like genes (DCLα and DCLβ), one exportin-5 gene (Exp5A), six argonaute genes (AGOA, AGOB, AGOC, AGOD, AGOE, and AGOF), and one RDR gene. These genes are conserved among oomycetes [25,45] and are homologs to those in phylogenetically distant organisms such as *A. thaliana*, *H. sapiens*, and *M. musculus*. These findings suggest that *P. capsici* may use canonical sRNA-mediated epigenetic regulation similar to that reported in mammals, plants, humans, insects, fungi, and other pathogenic oomycetes [82].

### Phylogenetic analysis and functional diversification of *P. capsici* sRNA regulatory enzymes

Phylogenetic analysis revealed that DCLα and DCLβ protein sequences clustered into two distinct groups, with DCLα related to the DCL1 group of oomycetes and DCLβ associated with DCL2 in *P. infestans*, *P. sojae*, and *P. ramorum*. This suggests that *P. capsici* DCL proteins may have a different evolutionary origin, as reported for DCL1 and DCL2 in several *Phytophthora* species [83], probably because of the functional divergence of the enzymes. For instance, the DCLα homolog in *P. infestans* is involved in the production of 21 nt sRNAs in the miRNA pathway [84], while the DCLβ homologs in *P. infestans*, *P. sojae*, and *P. ramorum* probably generate 25-nt sRNAs through the RNAi pathway [45]. However, further experiments are needed to clarify the roles of both enzymes in *P. capsici*.

Conversely, phylogenetic analysis of Exp5A from *P. capsici* indicated that it belongs to a diverse oomycete-specific clade, distinct from mammalian exportin-5 proteins. This suggests evolutionary divergence and potential functional specialization in the gene family. Notably, this is the first report of the presence and expression of an exportin-5 homolog (Exp5A) in *P. capsici* and other oomycetes; therefore, the evolutionary origin and function of oomycete Exp5A remain unknown. However, in mammals, exportin-5 is a protein in the caryopherin family responsible for transporting miRNAs through the nuclear pore from the nucleus to the cytoplasm in the presence of the Ran-GTP4 cofactor [82].

Interestingly, *P. capsici* encodes six AGO genes, while *P. infestans*, *P. ramorum*, *P. sojae*, and *P. parasitica* encode five, six, nine, and five AGOs, respectively. The AGOs of *P. capsici* contain domains (ArgoL, ArgoN, MID, PAZ, and PIWI) commonly found in functional Argonaute proteins [85–88]. Phylogenetic analysis grouped AGO1 from different oomycetes into clade I, whereas the other five enzymes (AGOB–AGOF) clustered into the more diverse clade II. Clade I AGOs may be miRNA-specific enzymes, whereas clade II AGO proteins may primarily process siRNAs [26,89]. Similarly, AGOC and AGOD of *P. capsici* clustered into separate clades, indicating paralogous divergence. However, based on our expression results, AGOC may be non-functional or conditionally regulated, as significant relative expression was observed only for AGOD.

In contrast, AGOB and AGOE were associated with AGOs from different oomycetes, whereas AGOF was associated with a distinct or novel clade, suggesting potential new functional roles for oomycete AGOs. These phylogenetic relationships of AGO proteins suggest gene expansion and possible functional specialization in sRNA regulatory pathways in *Phytophthora* species, which warrants further investigation. Finally, regarding the RDR proteins, our results show that *P. capsici* encodes a single RDR gene in its genome, similar to other oomycetes such as P. infestans, *P. sojae*, and *P. ramorum* [25]. In contrast, the phytopathogenic fungus *Verticillium dahliae* has been reported to encode three RDRs, whereas *A. thaliana* has six RDR homologs [92], suggesting that RDRs may have specific functions in different organisms and have distinct evolutionary origins, as well as unique gene regulation pathways among living organisms. Notably, the RDR gene in *P. capsici* is phylogenetically distant from other oomycete RDRs, indicating sequence differences between oomycetes. However, it remains unclear whether these differences lead to functional specificity. Based on the functions reported for members of the RDR enzyme family, it is possible that, similar to mammals, where RDR is involved in the biogenesis of secondary siRNAs from AGO-trimmed mRNA fragments [90,91], *P. capsici* RDR enhances gene silencing via AGO.

Notably, while most *Phytophthora* species encode a single RDR, many of these, reported for the first time in this study (i.e., *P. capsici*, *P. megakarya*, *P. cinammomi*, *P. rubi*, and *P. nicotianae*), expand the known distribution of RDRs within this genus. However, the genomes of *P. kernoviae* and *P. palmivora* do not encode RDR homologs. This could be due to a combination of functional reduction, replacement by other regulatory pathways, specific evolutionary pressures, evolutionary loss due to redundancy, and lower exposure to transposons or viruses, among other factors [108]. Identifying key enzymes in the sRNA pathway in *P. capsici* opens the door to further studies analyzing their functions to gain a deeper understanding of their roles in the lifestyle of the pathogen.

### Synteny, genomic organization, and protein-protein interaction networks of *P. capsici* sRNA regulatory enzymes

The genes of *P. capsici* involved in key enzymes of the sRNA pathway share syntenic relationships with homologous genes in other oomycetes, and the proteins they encode may interact with other proteins within complex networks. The DCLα, DCLβ, Exp5A, AGOA, AGOB, and RDR genes of *P. capsici* exhibited collinearity and were syntenic with the corresponding regions in the genomes of *P. infestans*, *P. sojae*, and *P. ramorum*, indicating conserved regulatory loci. A syntenic relationship has been reported for argonaute genes from *A. thaliana* and pineapple [93], as well as for some oomycete AGO genes [25]. However, for oomycetes, co-localization and syntenic conservation have only been reported for Ago3, Ago4, and Ago5 among *P. infestans*, *P. sojae*, and *P. ramorum* [25]. Synteny analyses between the genomes of *P. infestans*, *P. sojae*, *P. ramorum*, and *P. betacei* have revealed regions of high synteny in core genes and high plasticity in host-pathogen interaction genes, as well as segmental duplication events [109, 110]. Similarly, it has been observed that NRL genes *of P. infestans* are syntenic across the genomes of potato, *Arabidopsis*, tomato, and rice [111]. Therefore, our findings expand the understanding of conserved syntenic gene families involved in sRNA epigenetic regulatory mechanisms in oomycetes.

Regarding interaction networks, the results suggest that DCLα, DCLβ, Exp5A, and RDR proteins may participate in the interconnected silencing regulatory pathways. Different DCL-AGO-RDR combinations work synergistically to silence specific RNAs that control invading nucleic acids of either endogenous or exogenous origin, a process mediated by various sRNAs [94,95]. Interestingly, Exp5A was found to possibly interact with the Ran/TC4/GSP1 GTPase protein and various nuclear pore proteins, possibly indicating a transport mechanism similar to the yeast exportin system [96].

#### Host-dependent expression of *P. capsici* sRNA-related genes

There was a differential expression of sRNA-related genes in *P. capsici* isolated from infected chili pepper and broccoli leaves. Generally, chili pepper was more susceptible to infection by the pathogen than broccoli in the first infection, showing necrotic regions around the inoculation sites. These results are consistent with previous findings showing differences in aggressiveness levels across various hosts [81, 97, 98]. Moreover, *P. capsici* was isolated from broccoli leaves far from the inoculation site, which showed signs of infection in the form of small necrotic dots and decreased fluorescence emission. This indicates that, under controlled laboratory conditions, the pathogen can slowly infect broccoli leaves, a plant species often considered a non-host of *P. capsici* [99].

Our study showed that the central enzymes involved in the biogenesis, transport, and processing of sRNAs were significantly expressed in *P. capsici*. Specifically, the analysis of DCLs, AGOs, Exp5A, and RDR expression in *P. capsici* mycelium suggests that these enzymes are potentially functional in D3, Pc-chili, Pc-broccoli, Pc-ch-ch, and Pc-br-ch. However, host and infection history significantly altered the expression of these genes in both primary and secondary *P. capsici* isolates, which could influence gene regulation. Experiments on the DCL1 and DCL2 genes in *P. infestans* [100] and *P. sojae* [83] showed no significant differences in relative expression levels, supporting the idea that host and infection history in *P. capsici* affect the expression of DCLα and DCLβ genes.

The results of the relative expression of argonaute genes (AGOA, B, D, E, and F) suggest that they could play important roles in *P. capsici*, as shown by significant changes in their expression between treatments and the control, suggesting that they have distinct functions. Variations in the transcript levels of different AGO genes have also been observed in the mycelia of *P. sojae* and *P. infestans* [83,89], further emphasizing the differences in the potential functions of AGO genes in oomycetes.

Meanwhile, in *P. parasitica*, there is strong evidence that AGO3 plays a key role in virulence [26]. Therefore, we believe that the changes in the expression levels of different AGOs in *P. capsici* could be partly linked to the changes in pathogenicity observed during infection experiments, as potential sRNAs processed from the different AGOs might regulate effector genes [35], with the plant species and infection history also affecting the virulence of *P. capsici*. However, when the oomycete is recovered from infecting a first or mixed host, such as chili pepper and broccoli, it generally results in decreased gene expression compared to the control, with some exceptions. This suggests that oomycete gene activity increases due to the decreased expression of argonaute genes.

Finally, we observed variation in RDR expression across the studied treatments, indicating that broccoli infection significantly increased RDR expression compared to the control D3, Pc-chili, Pc-ch-ch, and Pc-br-ch. The importance of RDR expression in the vegetative mycelium of *P. sojae* has also been examined [83], revealing distinct sRNA-mediated silencing pathways within the genus *Phytophthora*.

#### Hypothetical model of *P. capsici* sRNA pathway

Based on data on the predictive subcellular localization, expression profile, and related literature on sRNA biogenesis and processing enzymes in oomycetes [25,26,89,101], we constructed a hypothetical model of the biogenesis, transport, and processing of sRNAs in *P. capsici* influenced by DCLs, Exp5A, AGOs, and RDR (Fig 7). This model includes five main steps: 1) Biogenesis of small RNAs in the nucleus and mitochondria by DCLα and DCLβ, respectively, from transcribed DNA (whether DCLβ activity occurs in the nucleus or mitochondria needs to be clarified); 2) sRNAs could be transported to the cytoplasm by exportin-5A and potentially recognized and loaded by one of the argonaute enzymes (AGOA, B, D, E, F) and carried out; 3) Processing of mature sRNAs; 4) Regulation of target genes; and 5) RNA-dependent RNA polymerase (RDR) synthesizes dsRNA from target gene residues and transports them via an unknown system to DCL enzymes to amplify the regulatory signal of a specific target gene.

**Fig 7.**
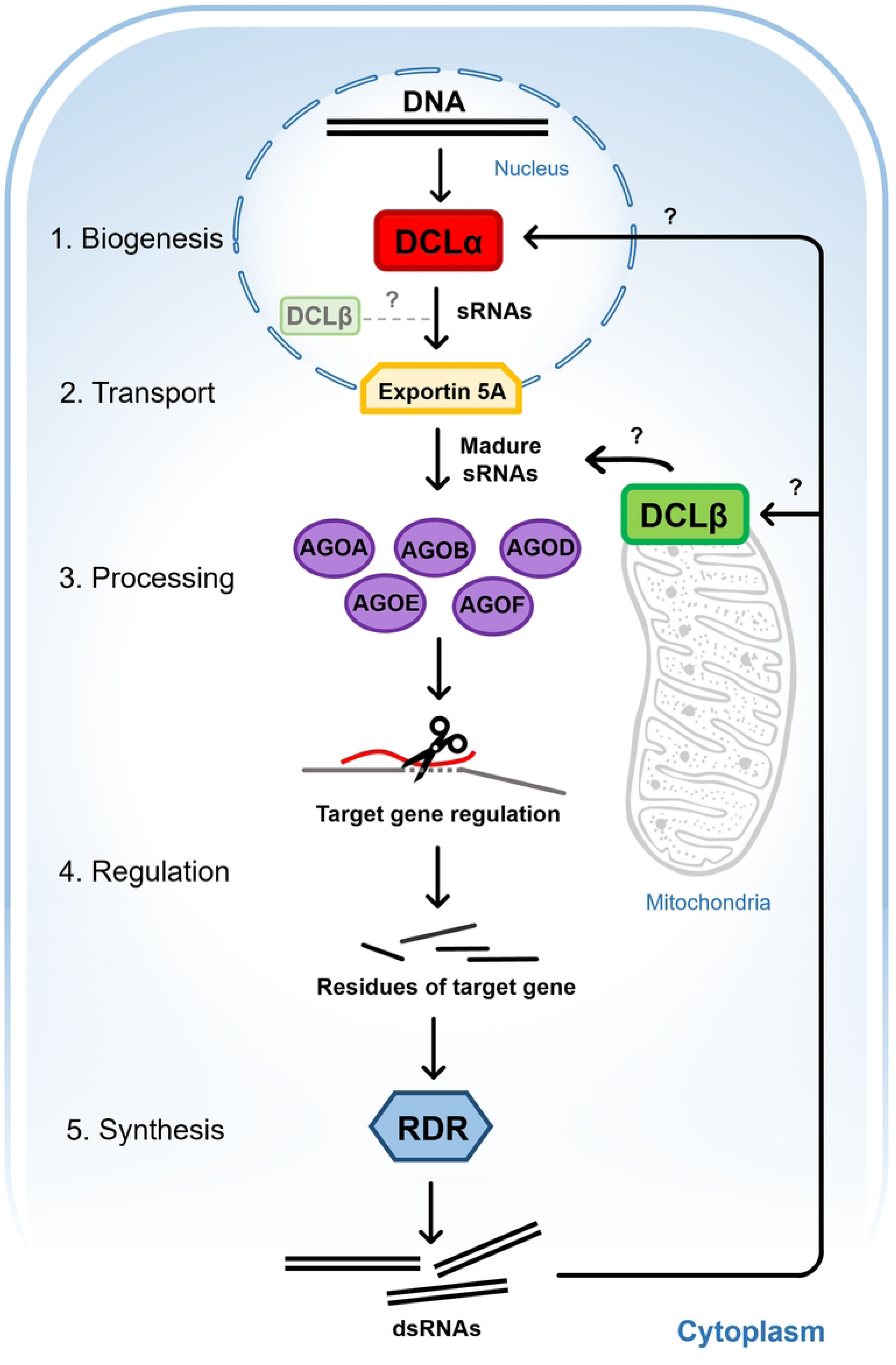
Hypothetical model of biogenesis, transport, and processing of small RNAs in *P. capsici.* 1. Biogenesis of small RNAs by DCLα and DCLβ, 2. transport of small RNAs by Exp5A, 3. Processing of mature sRNAs, 4. Regulation of target genes by AGOs, 5. Synthesis of dsRNAs by RDR to amplify the gene silencing signal.

## Conclusion

Our study offers the first comprehensive identification and expression profiling of core sRNA pathway components in *P. capsici*, an oomycete responsible for devastating various crops of agricultural importance worldwide. The *P. capsici* genome encodes essential enzymes involved in the small RNA pathway, including two DCLs, one Exp5A gene, six AGOs, and one RDR gene, which are phylogenetically related to homologous proteins from other oomycetes. Some of these genes are syntenic with homologous genes from other oomycetes. The presence and expression of these genes, especially Exp5A and RDR, which have not been previously reported in this species, highlight their potential roles in gene regulation during infection. Notably, we observed host-specific changes in sRNA gene expression, which may reflect transcriptional plasticity that contributes to the virulence and adaptation of *P. capsici* to both natural and non-host plants. However, the underlying mechanisms of epigenetic regulation mediated by these genes are complex and require further investigation, particularly to confirm functional diversification within different gene families. Finally, the present research expands the knowledge of epigenetic regulation mediated by sRNAs in *P. capsici*, paving the way for new strategies to control this pathogen and providing a foundation for future functional studies of RNA silencing machinery in oomycetes.

## Acknowledgments

Funding for this project was provided by PAPIIT-DGAPA-UNAM (Project #IN218024). We thank Secretaria de Ciencia, Humanidades, Tecnología e Innovación (SECIHTI) for a doctoral scholarship to JS-S and a postdoctoral scholarship to FUR-R by “Estancias posdoctorales por México 2022(1)”.

## Supporting information

**S1 Table. ID proteins phylogenetic reconstruction.** Sequence of Protein sequences and IDs used for phylogenetic reconstruction of DCL, Exportin-5, RDR and AGO.

**S1 Fig. Genomic localization of key genes in pathway of small RNAs.** Genomic localization of argonaut, dicer-like, exportin-5A, RNA-dependent RNA polymerase genes in *Phytophthora capsici.* Black lines indicate genes, blue: repeats.

**S2 Table. Locus synteny of key genes in pathway of small RNAs.** Locus synteny of DCLα, DCLβ, Exp5A, AGOA, AGOB, AGOC, AGOD, AGOE, AGOF and RDR of *P. capsici* versus *P. infestans, P. sojae* and *P. ramorum genomes.* X indicate No synteny.

**S3 Table Primers RT-qPCR.** Characteristics of primers and experimental conditions used to perform RT-qPCR of genes DCLα, DCLβ, Exp5A, AGOA, AGOB, AGOC, AGOD, AGOE, AGOF and RDR in *P. capsici*.

**S4 Table Ct value.** Ct value of DCLα, DCLβ, Exp5A, AGOA, AGOB, AGOD, AGOE, AGOF, RDR and EF-1α genes of *P. capsici* in RT-qPCR.

